# Lek-associated movement of a putative Ebolavirus reservoir, the Hammer-headed fruit bat (*Hypsignathus monstrosus*), in northern Republic of Congo

**DOI:** 10.1101/694687

**Authors:** Sarah H. Olson, Gerard Bounga, Alain Ondzie, Trent Bushmaker, Stephanie N. Seifert, Eeva Kuisma, Dylan W. Taylor, Vincent J. Munster, Chris Walzer

## Abstract

The biology and ecology of Africa’s largest fruit bat remains largely understudied and enigmatic despite at least two highly unusual attributes. The acoustic lek mating behavior of the hammer-headed bat (*Hypsignathus monstrosus*) in the Congo basin was first described in the 1970s. Then in the 2000s, molecular testing implicated this species and other fruit bats as potential reservoir hosts for Ebola virus and it was one of only two fruit bat species epidemiologically linked to the 2008 Luebo, Democratic Republic of Congo, Ebola outbreak. Here we share findings from the first pilot study of hammer-headed bat movement using GPS tracking and accelerometry units and a small preceding radio-tracking trial at an apparent lekking site. The radio-tracking revealed adult males had high rates of nightly visitation to the site compared to females (only one visit) and that two of six females day-roosted ∼100 m west of Libonga, the nearest village that is ∼1.6 km southwest. Four months later, in mid-April 2018, five individual bats, comprised of four males and one female, were tracked from two to 306 days, collecting from 67 to 1022 GPS locations. As measured by mean distance to the site and proportion of nightly GPS locations within 1 km of the site (percent visitation), the males were much more closely associated with the site (mean distance 1.4 km; 51% visitation), than the female (mean 5.5 km; 2.2% visitation). Despite the small sample size, our tracking evidence supports our original characterization of the site as a lek, and the lek itself is much more central to male than female movement. Moreover, our pilot demonstrates the technical feasibility of executing future studies on hammer-headed bats that will help fill problematic knowledge gaps about zoonotic spillover risks and the conservation needs of fruit bats across the continent.

## Introduction

Lek mating behavior is a very rare mating strategy among the more than 1200 known species of bats [1]. In the early 1970s it was the strong suspicion of lek behavior that led to Dr. Jack Bradbury’s interest in *Hypsignathus monstrosus*, or the hammer-headed bat, and he conducted the first detailed observational, acoustic, growth-rate, and radio-tracking movement study of *H. monstrosus* at a research station near Makoko, Gabon [2]. He concluded the population there used a classical lek mating system, as defined by a narrow set of distinguishing characteristics, as well as other traits commonly found among lekking species. In brief the ‘classical’ criteria of a lek species included, “(1) no male parental care, (2) an arena to which females come and on which most of the mating occurs, (3) the display sites of males contain no significant resources required by females except the males themselves, and (4) the female has an opportunity to select a mate once she visits the arena” [3]. Curiously, in West Africa this same species is reported to use a harem-based reproductive strategy suggesting the potential for intra-species behavioral plasticity (Dr. Dina Dechmann personal communication).

In Central Africa, the visitation of demographically mixed *H. monstrosus* at cacophonous lekking sites is an opportunity to reliably sample large numbers of individuals. Importantly, this species was epidemiologically linked to a human outbreak of Ebola virus (Luebo, Democratic Republic of Congo) [4] and has had repeated positive detections for *Zaire ebolavirus* antibodies and RNA [4–7]. An inhabitant of Central and West Africa, its geographical distribution overlaps with previous Ebola virus outbreaks within Africa [8]. Such aggregations of bats, at breeding or feeding sites, and their movements in between are important targets for epidemiological study as they may be key in determining spillover risk by increasing contact rates among individuals. For example, a biannual birth pulse with concurrent changes in immune response would introduce many susceptible individuals into a population and create a spillover pathway [9,10].

There is one study that reported visual observations of a month-long seasonal migratory movement of *H. monstrosus* from the Congo river upstream to the Lulua river in the Democratic Republic of Congo (DRC) [4], but little else is known about where these bats go and when. Beyond aggregations, movement data is critical to monitor animal responses to environmental change and to understand bat-human interfaces. In Australia, movement data helped reveal foraging shifts of flying-foxes from native (in decline) to non-native plants that may be bringing bats into closer contact with horses and human populations [11]. Furthermore, a few studies have suggested correlative associations between logging of fruit bat habitat and Ebola virus disease outbreaks, however, the findings to date remain inconclusive due to data limitations [12–15].

Here we present an analysis of hammer-headed bat movement in northern Republic of Congo from two pilot studies. The purpose of the pilots was to begin elucidating the movement ecology of a population of hammer-headed bats already the focus of a longitudinal virological study that began in 2011. That study includes monthly tree phenology observations that began in 2016 and local daily rainfall and temperature data collections since 2015. The distinctive honking sounds made by the congregations of males and the amount of bat activity during the initial and subsequent missions led us to believe the site is a lek [2]. And upon listening to calls recorded and digitized by the Macaulay Library, we recognized that we often hear the ‘staccato buzz’ performances, which Dr. Bradbury observed to occur when a female passes within meters of a male, for example, *Hypsignathus monstrosus* (<u>ML170632</u>). We began with an affordable VHF radio-tracking study that helped inform the selection of a base station methodology that we then used for the GPS-based second pilot. With data collections completed, here we describe our findings and revisit Dr. Bradbury’s definition of a lekking species to support or reject our initial hypothesis of the site as lek. As pilot efforts our sample sizes are low but our findings begin to reveal some insights on the unique lek-associated movement ecology of male and female hammer-headed bats and present future opportunities to characterize bat-human spillover interfaces. We also aim to help inform and improve future movement ecology studies focused on this species.

## Material and methods

### Radio-tracking deployment December 2017

We conducted a preliminary tracking effort in December 2017, using basic VHF trackers (Holohil BD-2, 0.75 g). Following capture and sample collection, 10 bats (4 adult males, 4 adult females, and 2 juvenile females) were hand restrained and 100% ethyl isocyanoacrylate glue, which cured within minutes, was used to attach the trackers. Hair was layered over to improve the attachment. Nine of the bats were released and monitored for 10 days and one adult male was monitored for nine days.

### GPS unit deployment April 2018

In April 2018, we tagged 11 bats with Bird Solar 15 g units (E-obs Digital Telemetry) programmed to collect a burst of five GPS points every 30 minutes and accelerometry measurements every 10 min on three axes in the evening between approximately 18h00 West Africa Time (WAT, local time) and 06h00 WAT. A subset of the heaviest male and female bats were tagged beginning on April 12 and ending April 21.

The Bird Solar 15 g units were attached to a pre-made ‘capes’ cut from soft and pliable six ounce mouse grey fabric (Dimension Polyant Fabrics - X-Pac™ VX21) to fit neck circumferences of 100-125 mm in ∼5 mm increments (Fig 1). A 3/8” plastic side-releasable buckle was attached to the cape with Seam Grip (Gear Aid™) and cured 24 hrs in a clamp. Fabric areas that received Seam Grip were pre-treated with 99% alcohol and abrasion to remove the fabric’s durable water-repellent finish that could reduce bond strength. On site with hand restraint, capes were custom-fit to each individual, allowing enough room for a handler to insert a finger between the collar and neck but small enough to attempt to prevent the collar from slipping over the head and ears. Once fit, the cape was removed and the bat was placed in a cloth bag. The unit was glued to the cape with a mix of UV Aquaseal and Cotol-240™ (to accelerate curing to time to ∼2 hours) along with clamping. After curing the unit was labeled with a recovery phone number and then reattached to the bat with the buckle. Total cape and unit mass was 18 g.

**Fig 1.**
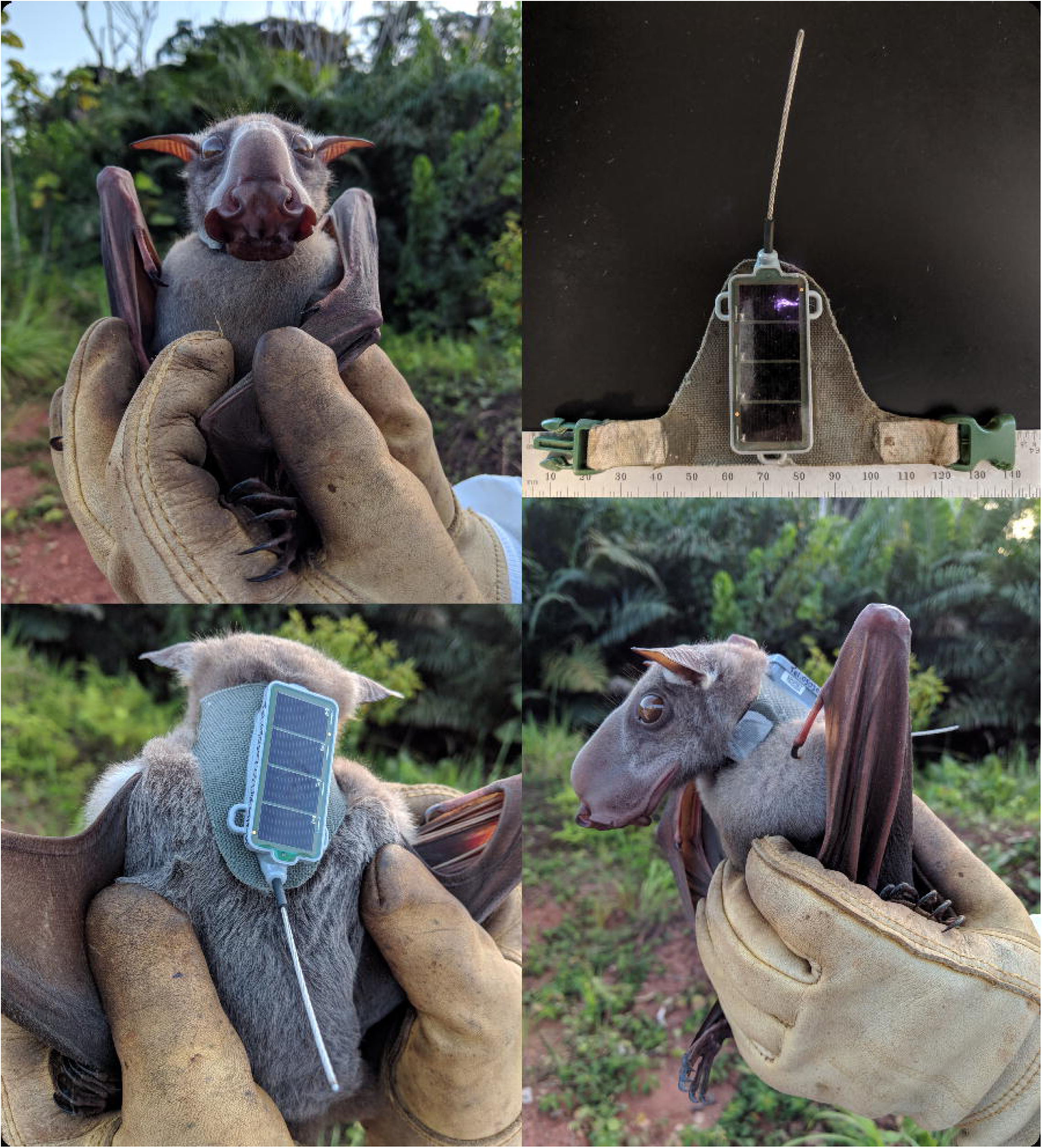
Adult male hammer-headed bat with solar GPS tag and collar harness.

Initially both collared and control bats were held from 12 to 24 hours (depending on initial capture time) for observation in an isolated and shaded mesh cage (5.0 m × 5.0 m × 1.8 m). The cage (flexible poultry netting covered partially with leaves) permitted short distance flight and was enriched with large leaves for cover, roosting branches, as well as food and water ad libitum. Care was taken to avoid disturbing bats in the cage and behavioral observations were made from a distance. Bats were initially clearly stressed by the cage and flew repeatedly into the mesh walls of the cage, apparently unaware it was there, until they established a roost on the cage roof. Beyond the cage stress, we observed no adverse impacts of the collar on behaviors in the initial eight bats that were collared and three bats that were included as controls. As no adverse collar issues were noted and substantial cage stress was observed we opted to release the last three bats without any cage observation period.

Data from the units were remotely downloaded from each individual bat when in proximity of a base station (BaseStation II, E-obs Digital Telemetry). We fixed a permanent base station with a large antenna (Omni-Antenna 868MHz 3dBd) in a small clearing used for netting the bats in the lek. Mid-mission we temporarily placed the fixed base station and large antenna on a vehicle that drove at 5 km/hr and surveyed Route N2 15 km northeast and southwest of the site. In addition, during the 10 day deployment mission we patrolled Route N2 ∼3 km southwest or northeast of the lek. Then each day through May 28, 2018, we used a hand-held base station to survey along the road between the lek and the southern extent of Libonga village. Between 14– 16 February, 2019, the team macheted their way through the forest to visit male and female frequently used day-roosts with a hand-held base station. A return visit to the female’s primary day roost was made on 7 June, 2019.

For analysis, we selected one of the five GPS burst points so that each reported ‘GPS location’ was spaced approximately 30 min apart. Individuals with 24 hours or less of movement (i.e from the night of release) were excluded from further analyses. We classified flying and non-flying behavior from the accelerometry data according to Abedi-Lartey et al. [16]. We transformed latitude and longitude locations from a WGS 84 coordinate system to UTM coordinates (UTM zone 33N) to calculate distances in meters and calculated the nightly proportion of GPS locations that were within 1 km of the lek (0.91358°N, 15.59132°W) as a measure of site visitation. Only nights with more than 10 GPS locations were used for calculations of nightly distance traveled, which excluded 13 bat nights.

To visualize bat movement between 12 April and 31 May, we created a movie using the R package moveVis (S1 File). The utilization distribution of each bat was mapped using a classic kernel density estimation (KDE) for each bat with vertices set to 80% and 95% location probability [17] (R package: adehabitatHR). Statistical tests and mapping were conducted in R v3.5.2 [18]. All animal handling, tracking, and capturing activities were approved under Wildlife Conservation Society IACUC #17-05, NIH ASPs #2015-010 and #2018-015-F, and by Republic of Congo Institut National de Recherche Forestière (IRF) Permit #260. The GPS unit data is publicly accessible on Movebank (www.movebank.org) under study name ‘Hypsignathus monstrosus Olson Central Africa’.

## Results

### Radio-tracking effort December 2017

All male detections occurred at our lek site (S2 File). After release, two of the adult males were detected at the lek for nine consecutive days (including day roosting at the site) and the two other males were detected there for one or three consecutive days (no day roosting). Only one female, an adult, was detected at the lek four nights after she was initially tagged. The day after release, this same female and a juvenile female were detected day-roosting 100 m west of Libonga village, located ∼1.6 km southwest from the lek (Fig 2). The same juvenile was found at that same day-roost for eight consecutive days. We determined that the tag VHF signal was detectable through the forest at distances up to 800 m. Further tracking efforts 3 km northeast and 3 km southwest along Route N2 were unsuccessful. The lack of easily navigable paths beyond the main road and the detection rates at the lek and village motivated us to pilot a GPS tracking unit with remote download, more suited to identify precise feeding and movement strategies of these bats.

**Fig 2.**
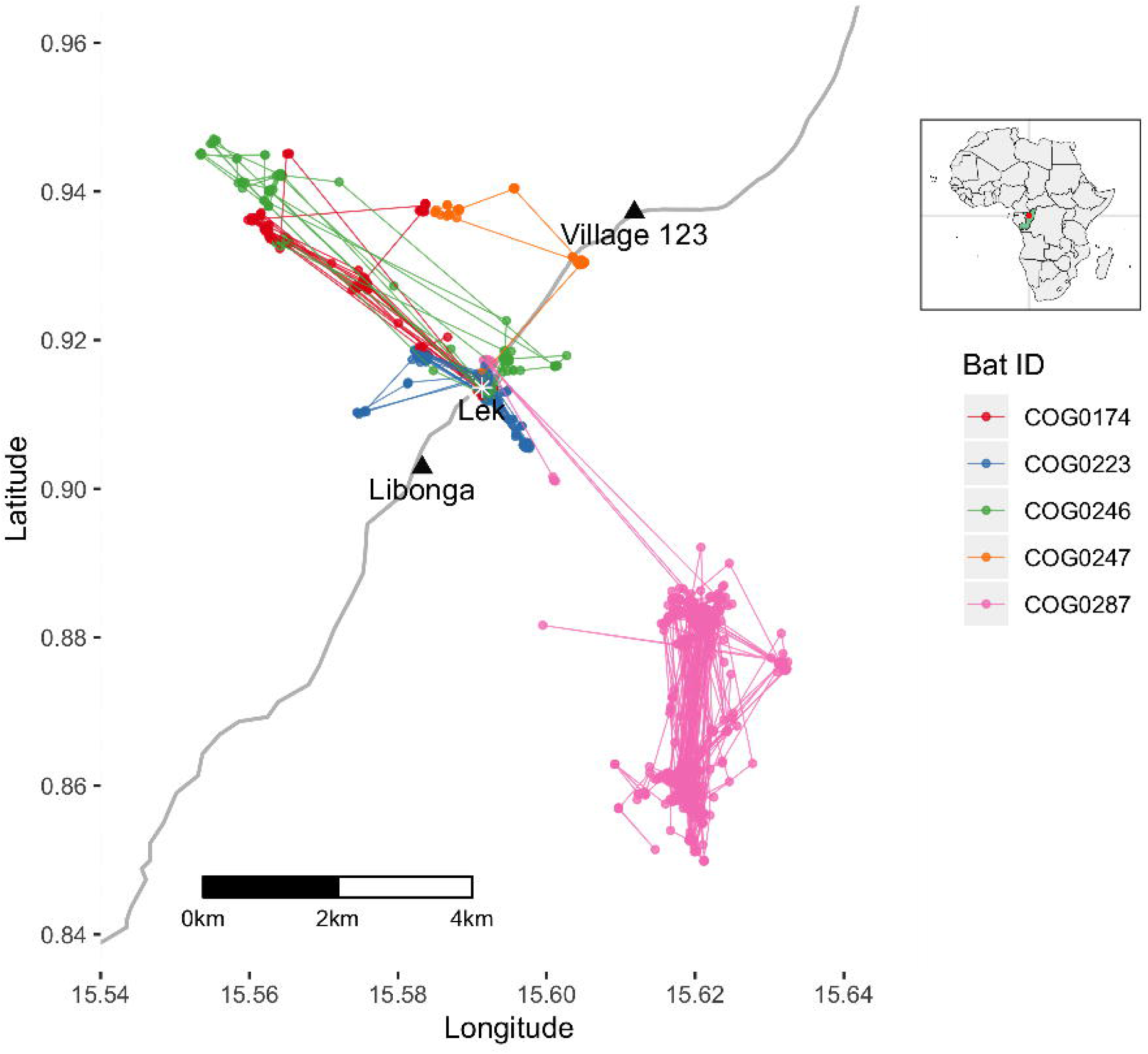
GPS tracking of five bats. Locations and movement and of bats beginning mid-April 2018. The main north-south paved N2 highway is shown in grey (inset map from R package: rworldmap). The estimated center of lek activity is located at the star and locations of the two nearby villages are shown (black triangles).

### GPS unit deployment April 2018

Eleven bats, comprised of four females and seven males, were collared and released (Table 1). We recovered the collars deployed on bats COG0222 and COG0246 with intact buckles and no indication of an associated carcass and concluded that these individuals managed to slip the collar off over their heads. One of the recovered collars was redeployed. In total five individuals provided tracking data while tracking ended prematurely on six individuals.

**Table 1.** Deployment summary for all tagged bats during the April 2018 study period. The table is sorted by the number of GPS locations and additional statistics are shown for the five bats with over 24 hours of GPS data.

| Bat ID | Sex | Weight (g) | Date deployed | Duration of GPS activity (days) | Number of GPS locations | Mean time to first fix (s) | Nights with fixes | Percent of fixes < 1 km of site | KDE 80% (ha) | KDE 95% (ha) |
| --- | --- | --- | --- | --- | --- | --- | --- | --- | --- | --- |
| COG0287 | f | 296 | 16-Apr | 305.5 | 1022 | 22.1 | 53 | 2.2% | 462 | 1010 |
| COG0223 | m | 405 | 13-Apr | 17.3 | 441 | 20.2 | 18 | 73% | 85.6 | 185 |
| COG0174 | m | 464 | 21-Apr | 7.0 | 177 | 21.0 | 8 | 24% | 971 | 1620 |
| COG0246 | m | 425 | 14-Apr | 6.5 | 173 | 20.0 | 7 | 40% | 614 | 1290 |
| COG0247 | m | 409 | 14-Apr | 2.4 | 67 | 25.5 | 3 | 7% | 439 | 829 |
| COG0222 | m | 406 | 13-Apr | 0.5 | 25 | 18.3 | 1 | - | - | - |
| COG0291 | f | 275 | 16-Apr | 0 | 4 | 25.8 | 2 | - | - | - |
| COG0206 | m | 445 | 12-Apr | 0 | 2 | 27.0 | 1 | - | - | - |
| COG0207 | f | 271 | 12-Apr | 0 | 2 | 43.2 | 1 | - | - | - |
| COG0248 | m | 406 | 14-Apr | 0 | 2 | 31.6 | 1 | - | - | - |
| COG0224 | f | 293 | 13-Apr | 0 | 1 | 51 | 1 | - | - | - |

Our lek was a more central feature of the movements and utilization distributions of male bats compared to the female bats (Fig 2). The lek location was within the 80% KDE of night location probability for all four males whereas only the female’s 95% KDE of night location probability included the site (S1 Fig, S1 Table). The female’s average displacement from the lek was 5504 m (range 379 m – 7789 m) and was significantly greater than locations for all males combined, at 1442 m (range 80 m – 5470 m; two-sample t(1633) = 60.3; p = 2.2e-16). For the males, 51% of their fixes were located within 1 km of the lek compared to only 2.2% of the female’s fixes (Chi-squared = 603.4; df = 1; p-value < 2.2e-16). Overall, the female’s mean daily distance traveled was 10636 m (range 3363 m – 17560 m) and was significantly greater than the mean daily distance traveled by all males combined, at 6088 m (range 1097 m – 18254 m; two-sample t(71) = 5.1, p = 0.000003).

On average these hammer-headed fruit bats showed fidelity to certain day-roost sites within a few hundred meters (Fig 3). They also typically reached their furthest distance from the day-roost site between midnight and 2h00 (S2 Fig). Home range size varied between 185 and 1010 ha based on 95% KDE (S1 Table).

**Fig 3.**
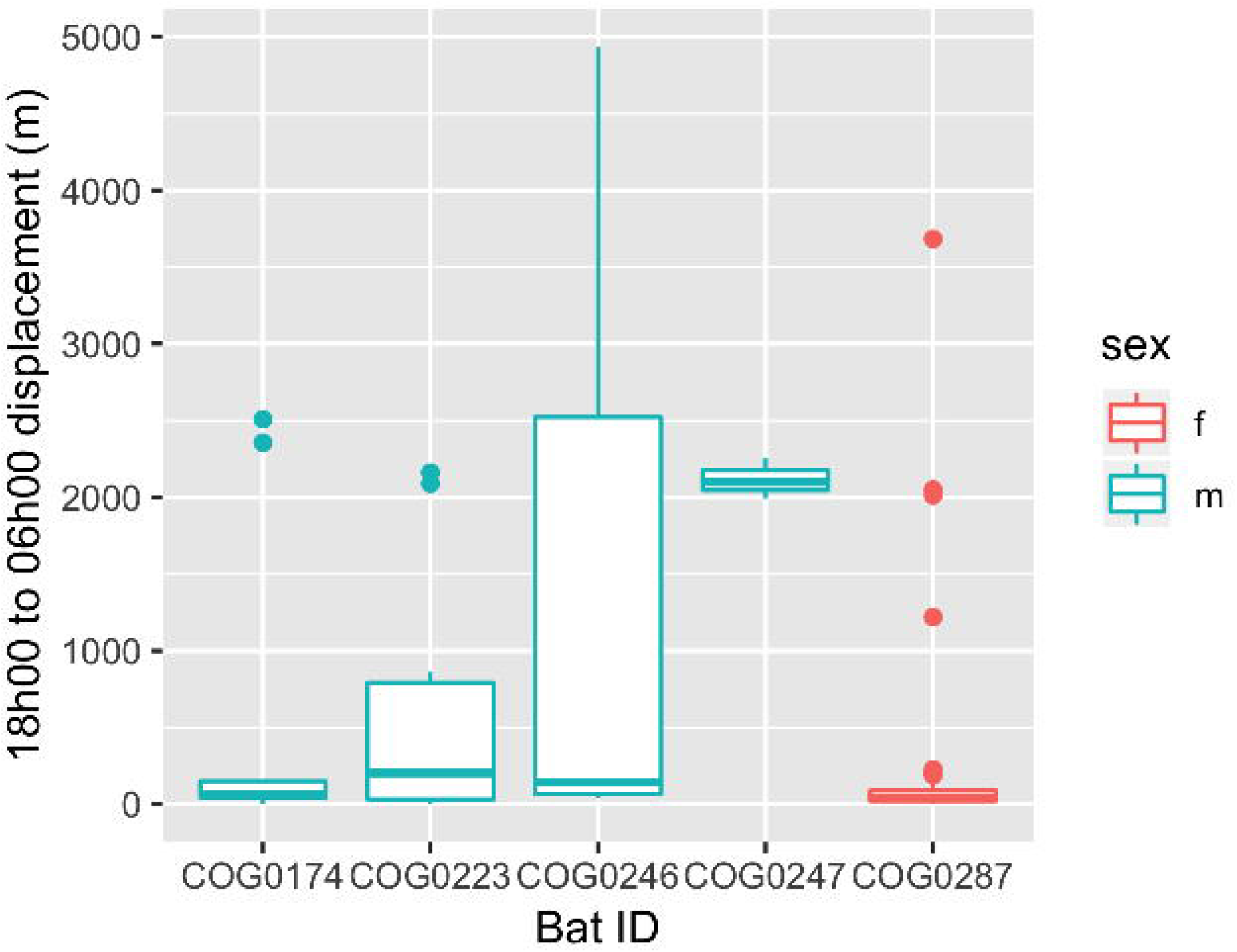
Day roost fidelity. Tukey box and whiskers plot of nightly overall displacement patterns (m) by each bat as measured from where the bat began the evening (typically around 18h00 WAT) and arrived in the following morning (typically around 06h00 WAT).

For the majority of the males, site visitation was daily. Three of the male bats visited the site every night, totaling 7, 8, and 18 consecutive nights for bats COG0246, COG0174, and COG0223 respectively. Although the GPS tracking data for the fourth male (COG0247) does not show any re-visitation to the site, the base station at the lek site logged communications and incomplete download attempts with COG0247 on April 20, 25, 26, and 28.

The female (COG0287) appeared to not visit the site for at least a period of 35 days from 21 April through 26 May, when 30 minute nightly GPS logging capability was lost due to low battery power. With the decline of tag power, the tag’s base station communication remains functional, but the tag decreases the number of GPS fixes and eventually shuts off GPS. The lek base station downloads revealed she visited the site in 2018 on 12 June, 24 July, 30 July, and 24 August. Though her two initial visits were detected by GPS and not the base station, a rough estimate of mean site visitation frequency is 22 days with standard deviation 22 days. All data from her unit was last downloaded on 16 February 2019 when the team detected her at an intensely used day roosting site (0.8821882°N, 15.61998°W).

Overall flight activity for both male and female bats, whether detected by displacement or accelerometry, appears greatest early in the evening and trends downwards until a spike in activity before dawn (Fig 4). Classification of the accelerometry data detected 9253 flying or not flying observations. During nights, 651 of the accelerometry data observations were classified as flying and 8147 were classified as ‘not flying’, a ratio of flight:resting observations of about 2:25. There were 455 ‘not flying’ accelerometry observations and no ‘flying’ observations during locally calculated daytime suggesting bats are likely stationary during daylight hours. Lastly, we observed the female’s nightly displacement was negatively correlated with lunar illumination (S3 Fig; adjusted r-squared = 0.168; intercept = 12898; slope = −4849; p-value = 0.004519), but there was no correlation of displacement and lunar illumination among the males (adjusted r-squared = −0.0148; intercept = 5574; slope = 1329; p-value = 0.480).

**Fig 4.**
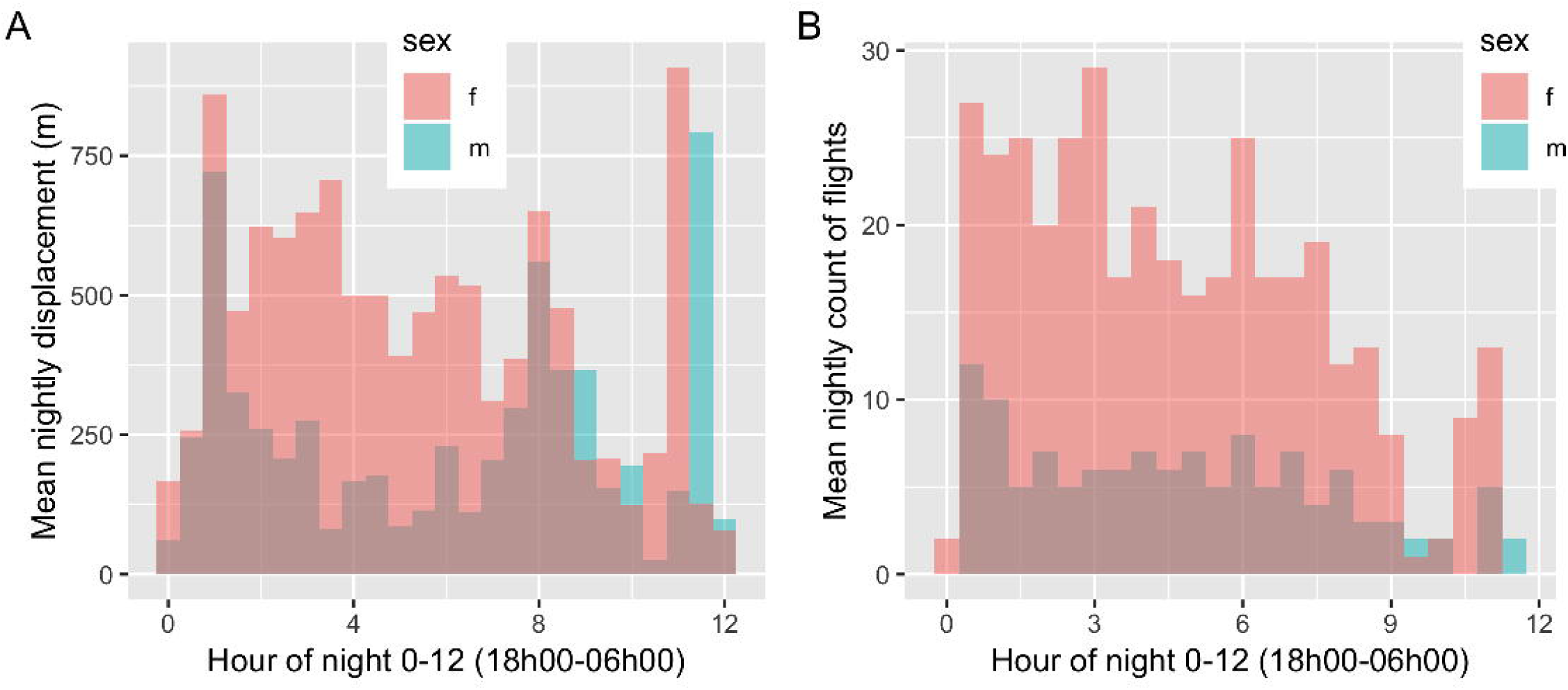
Nightly bat flight activity. (A) Barplot of GPS-based distance flown every 30 min beginning at 18h00 (time zero in plot) and 06h00 (time 12 in plot) and (B) of flight behavior observations as determined by accelerometry over the same period.

## Discussion

Our pilots provide a glimpse into hammer-headed bat movement ecology and further support our original assessment of the site as a lek for this fascinating species. The VHF and GPS pilots both indicated males frequently visit and occupy the site and that the site is visited, albeit much less frequently, by female bats. We found 51% of fixes for all males combined and 2.2% of the fixes for the female were within 1 km of the site and the latter’s 80% KDE showed no overlap with the site. Hence, we infer the lek does not contain ‘significant resources required by females except the males themselves’ [2]. While we are not yet able to visualize nightly bat activity in the canopy at the site, the regularly audible sound of ‘staccato buzzes’, which has been correlated with copulation, supports a conclusion that adult females are coming to our site for mating purposes [2]. Worth noting, the site is not exclusive to nonpregnant females as the majority of adult females captured for the virological study are pregnant. In addition, juveniles and subadults of both sexes are commonly sampled (authors’ unpublished data). We conjecture the lek visitations by non-reproductive individuals are associated with learning and social behaviors.

In general, movement patterns matched the magnitude of those observed by Bradbury’s radio-tracking study [2]. His population had ‘local’ day-to-day shifts on the order of 34-74 m, longer distance roost-shifts on the order of 5 km, and nightly flights up to 10 km. There is a previous report of hammer-headed bat migration at the southern extent of their range occurring in April and May [4,19]. Our study, located in the core of their range did not detect migratory behavior, but the male tracking only covered a maximum of 18 days and the female had reporting gaps of 42 and 53 days between April 2018 and February 2019.

Even though our unit performance was similar to other studies, interpretation of our findings should carefully consider our study limitations [20]. The tracking system relies on a receiver station to download data from the tags, and one station is fixed at the lek itself, so the majority of data collection is biased against bats that visit the lek frequently. We attempted to mitigate this bias by daily monitoring the female day roost identified during the VHF pilot, and by making a much delayed trek out to her day roost site in mid-February. We suspect our inability to detect the other females may have been due to their less frequent visits to the lek site and wider dispersal than males as well as collar loss (as we observed with two males). These data are consistent with recapture data at this site in which females are recaptured less frequently than males and with larger gaps between recaptures, suggesting that females are transient visitors to the lek site (Dr. Vincent Munster, personal communication). Based on the VHF data we had anticipated that a proportion of the females would be roosting near Libonga village. However, we were not logistically able to regularly visit sites off the main road and so a visit to the day roosting site of the female (COG0287) was delayed until February 2019. The GPS units were conservatively programmed to run 3-4 weeks but we were rather surprised to find the solar recharge on a nocturnal species roosting in the upper tree canopy was sufficient for a unit to log a location after 305 days.

Our findings will help future GPS-based movement ecology studies be better equipped and prepared to maximize their data collections for different research questions and tracking methodologies. This can help mitigate the already often high transaction costs of conducting research in remote corners of Africa. For example, we can now make more appropriate staffing and logistical plans for future deployments to visit the location of each bat’s regular day roost site. In addition, species-specific programming can improve and extend the performance of GPS units [20], and so we have subset the female’s tracking data to model how programming for fewer fixes (which would allow longer deployments) each night would influence her home range calculations. We are optimistic that the recent advancements in tracking technology, e.g. the <u>ICARUS Initiative</u> or <u>Motus</u>, combined with the growing number of bat movement studies can help support their conservation and protect human health.

## Supporting information

S1 Table. Utilization distributions.

S1 File. Movie of bat movement.

S2 File. VHF tracking observations.

## Acknowledgements

We are grateful to the local support teams behind each bat field mission, including camp cooks, support staff and the bat catching team. The effort of engineering a light weight collar greatly benefited from consultations with and donations of collar supplies from Graham Williams at Cilo Gear as well as Patrick Odenbeck and the team at Mystery Ranch. A special thanks also to Nistara Randhawa for graciously loaning us an E-obs tracker to help design the collar and to Kenneth Cameron for testing earlier designs. Marc Buentjen and Franz Kuemmeth at E-obs provided reliable and timely trouble-shooting and programming support. This work was supported by the Intramural Research Program of the National Institute of Allergy and Infectious Diseases, National Institutes of Health.

## Disclaimer

The contents are the responsibility of the authors and do not necessarily reflect the views of the National Institutes of Health, or the United States Government.

## Author Contributions

**Conceptualization:** Sarah H. Olson, Chris Walzer, Dylan W. Taylor, and Vincent J. Munster

**Data curation:** Sarah H. Olson

**Formal analysis:** Sarah H. Olson

**Funding acquisition:** Vincent J. Munster, Sarah H. Olson, Chris Walzer

**Investigation:** Sarah H. Olson, Gerard Bounga, Alain Ondzie, Trent Bushmaker, Stephanie N. Seifert, Chis Walzer, and Eeva Kuisma

**Methodology:** Sarah H. Olson, Chris Walzer

**Project administration:** Sarah H. Olson, and Eeva Kuisma

**Resources:** Sarah H. Olson, and Dylan W. Taylor

**Software:** Sarah H. Olson

**Supervision:** Sarah H. Olson, Alain Ondzie, Eeva Kuisma, and Chris Walzer

**Validation:** Sarah H. Olson

**Visualization:** Sarah H. Olson

**Writing – original draft:** Sarah H. Olson

**Writing – review & editing:** Sarah H. Olson, Dylan W. Taylor, Trent Bushmaker, Stephanie N. Seifert, and Chris Walzer

## Supporting information

**S1 Fig.**
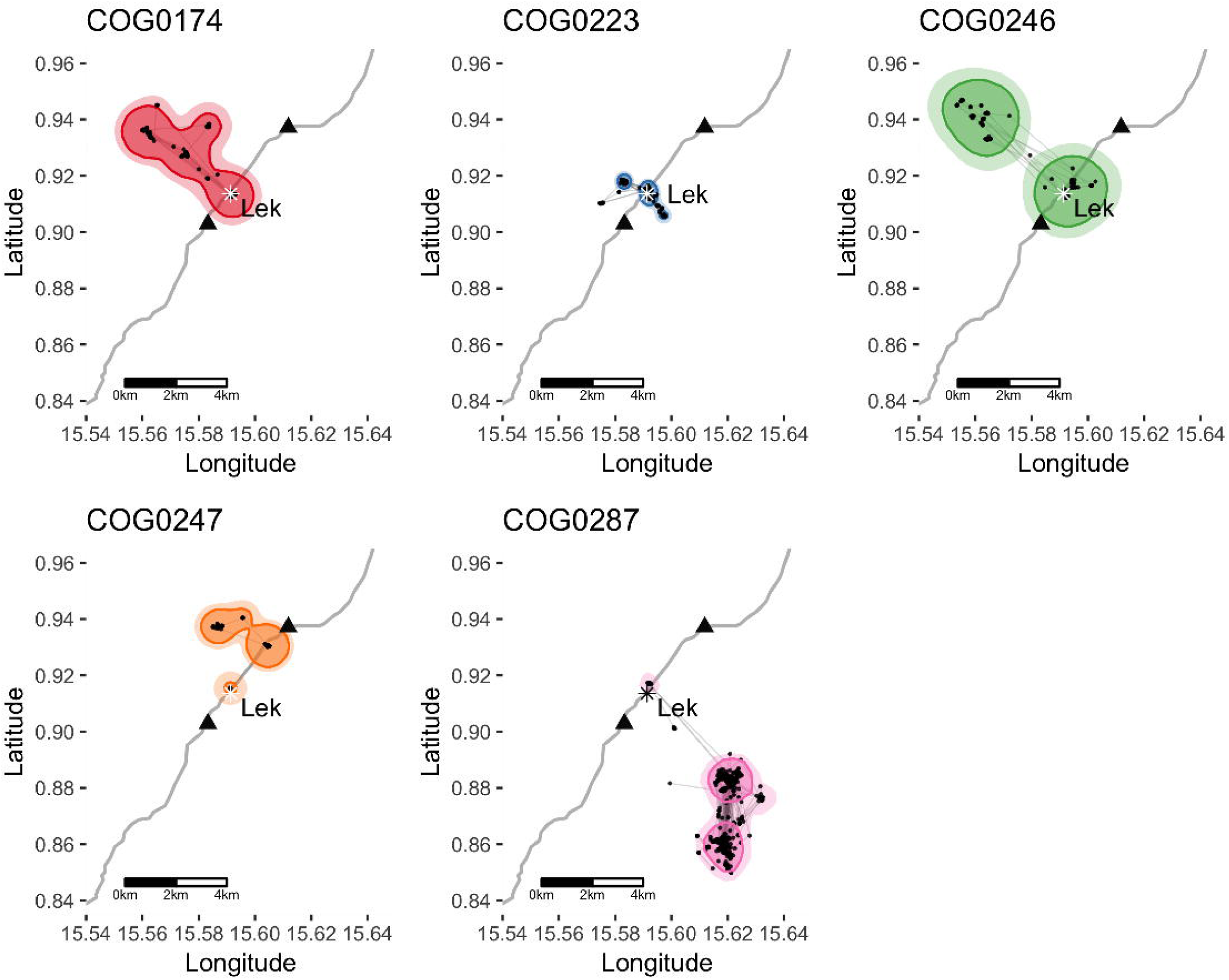
Kernel density estimation of each bat’s utilization distribution. Contours of 80 (darker with outline) and 95 (lighter) percent probability of utilization for each bat.

**S2 Fig.**
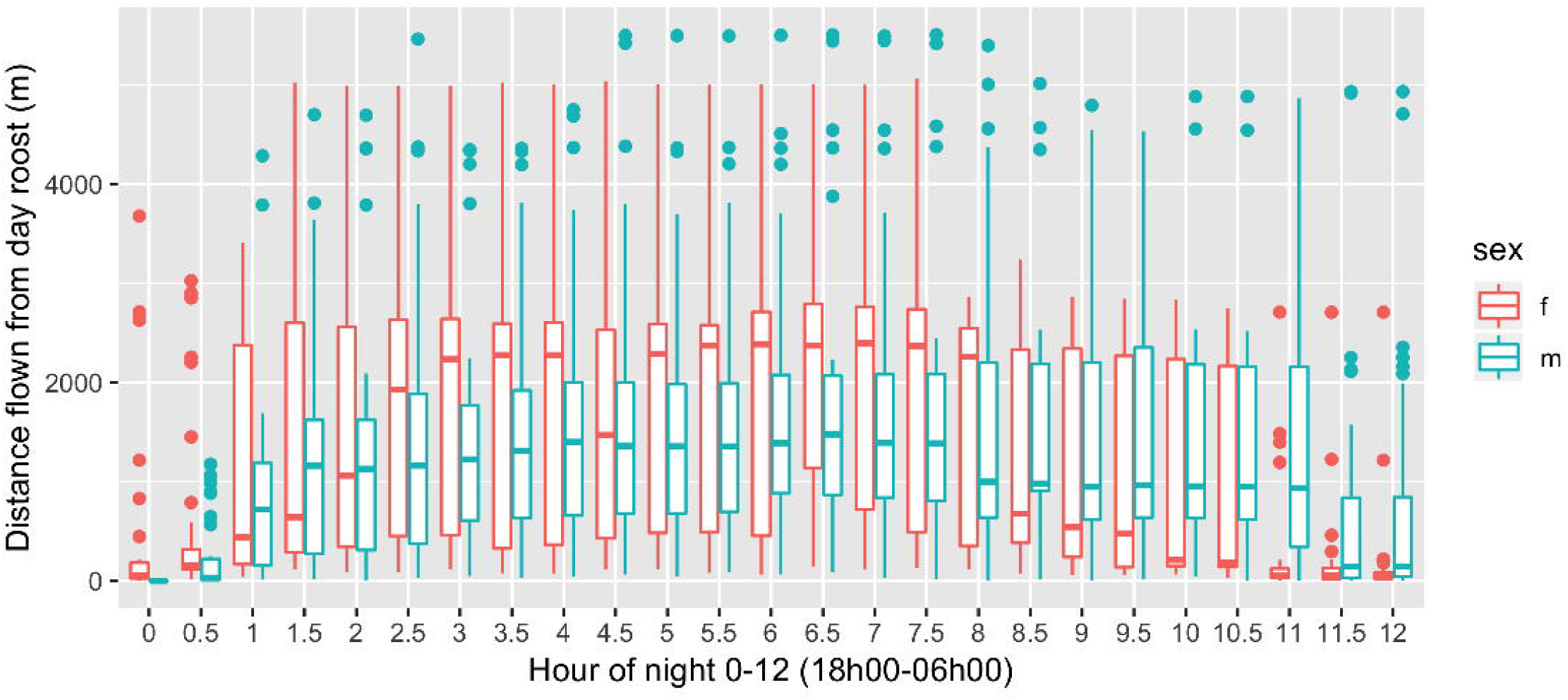
Tukey box and whiskers plot of displacement from day roost location each night. Day roost location is defined as the first GPS location of the evening typically around 18h00 (time 0 in the figure). Displacement is observed over 30 min intervals for all males combined and separately for the female.

**S3 Fig.**
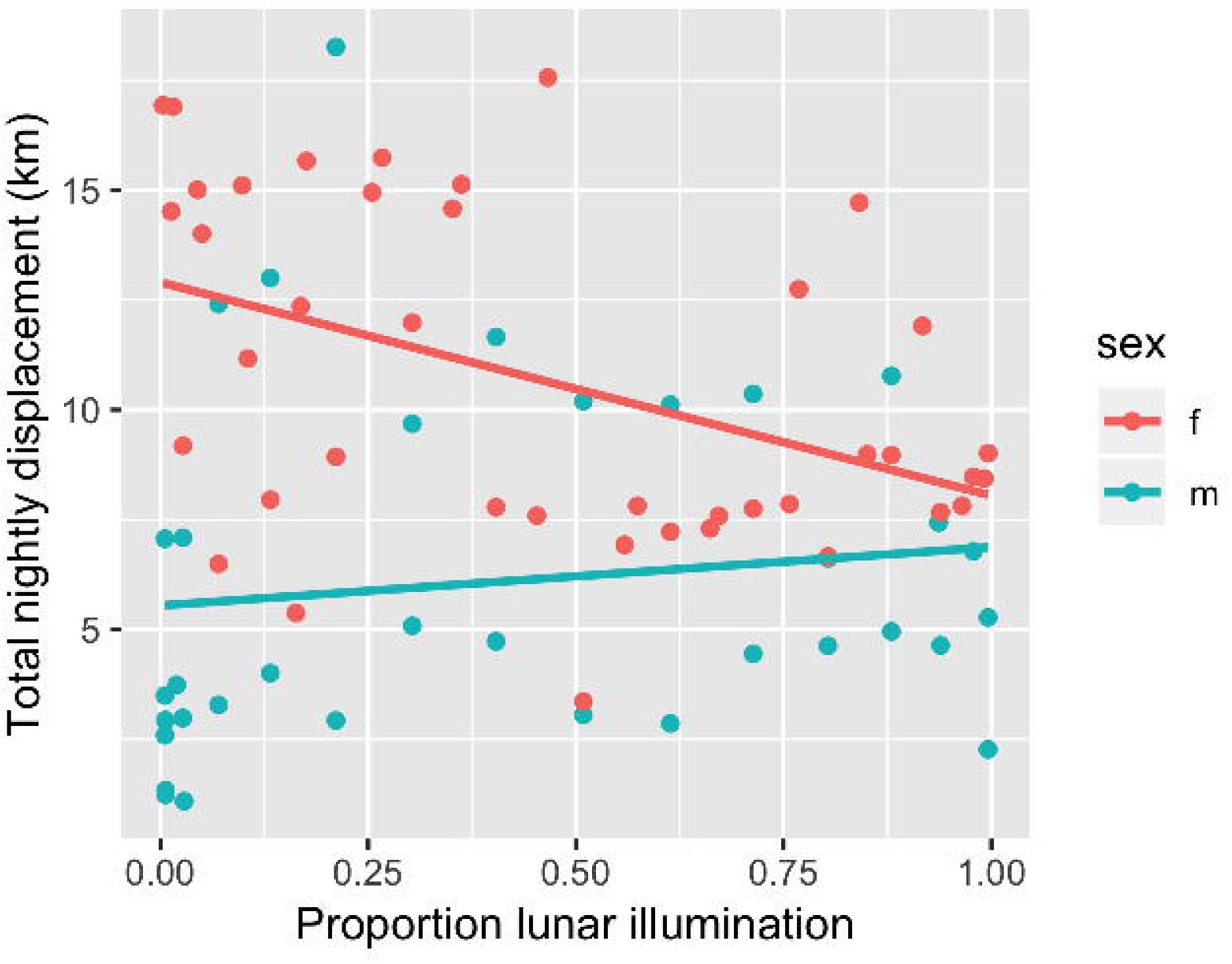
Plot of daily lunar intensity and nightly bat movement by sex. The female’s nightly displacement is negatively correlated with nightly lunar illumination (adjusted R2 = 0.168; intercept = 12898; slope = −4849; p-value = 0.004519). There is no correlation of displacement and daily lunar illumination among the males (adjusted R2 = −0.0148; intercept = 5574; slope = 1329; p-value = 0.480). Data excludes 13 bat nights when less than 10 GPS locations were collected.

**S1 File. Movie of bat movement.** The single female bat, COG0287, is tracked in pink. Each individual’s track fades after 24 hrs. Local time (WAT) is displayed. Route N2 is the faintly visible diagonal road along a southwest and northeast axis. Basemap data from Mapbox Satellite <u>© Mapbox</u>, <u>© OpenStreetMap</u>, and <u>© DigitalGlobe</u>; <u>improve this map</u>.

**S2 File. VHF tracking observations.**

**S1 Table.** Utilization distributions (ha) for individual bats calculated using minimum convex polygons (90%) and kernel density estimation (80% and 95%).

