## Supplementary material for "Lek-associated movement of a putative Ebolavirus reservoir, the Hammer-headed fruit bat (*Hypsignathus monstrosus*), in northern Republic of Congo": S1 Table. Utilization distributions.

**S1 Table.** Utilization distributions (ha) for individual bats calculated using minimum convex polygons (90%) and kernel density estimation (80% and 95%).

| Bat ID | Minimum convex polygon (90%) (ha) | Kernal density estimation (80%) (ha) | Kernal density estimation (95%) (ha) |
|---|---|---|---|
| COG0287 | 615.9 | 461.5 | 1010 |
| COG0223 | 108.5 | 85.6 | 185.3 |
| COG0174 | 547.3 | 970.6 | 1621 |
| COG0246 | 614.5 | 1293 | 2414 |
| COG0247 | 97.07 | 438.5 | 829.2 |
